# FebRNA: an automated fragment-ensemble-based model for building RNA 3D structures

**DOI:** 10.1101/2022.04.25.489348

**Authors:** Li Zhou, Xunxun Wang, Shixiong Yu, Ya-Lan Tan, Zhi-Jie Tan

**Author notes:** Corresponding authors (to ZJT); (to YLT). These authors contributed equally to this work (Li Zhou, Xunxun Wang).

## Abstract

Knowledge of RNA 3-dimensional (3D) structures is critical to understanding the important biological functions of RNAs. Although various structure prediction models have been developed, high accuracy of predicted RNA 3D structures is still limited to the RNAs with short length or with simple topology. In this work, we proposed a new model, namely FebRNA, for building RNA 3D structures through fragment assembly based on coarse-grained (CG) fragment ensembles. Specifically, FebRNA is composed of four processes: establishing the library of different types of CG fragment ensembles, building CG 3D structure ensemble through fragment assembly, identifying top-1 CG structure through a CG scoring function, and rebuilding the all-atom structure from the top-1 CG one. Extensive examination on different types of RNA structures indicates that FebRNA gives consistently reliable predictions on RNA 3D structures including pseudoknots, 3-way junction, 4-way and 5-way junctions, and RNAs in the RNA-Puzzles. FebRNA is available at website: https://github.com/Tan-group/FebRNA.

## INTRODUCTION

Noncoding RNAs have various important biological functions such as gene regulation and catalysis [1,2], and the functions of noncoding RNAs are generally related to their native structures, especially three-dimensional (3D) structures [3-12]. Therefore, the knowledge of RNA 3D structures is crucially helpful not only for understanding various functions of natural RNAs, but also for designing artificial RNAs and related drug molecules [3,8,13].

Experimental methods including X-ray crystallography, NMR spectroscopy and cryo-electron microscopy have been widely used to experimentally determine RNA 3D structures [14]. However, due to the high cost of experimental methods, the number of RNA structures deposited in protein data bank (PDB) is still limited until now. As complementary methods, various computational models have been developed to predict RNA 3D structures in silico. These models can be roughly classified into physics-based ones such as SimRNA [15,16], iFold [17], NAST [18], IsRNA [19,20], HireRNA [21], oxRNA [22], and our model with salt effect [23-26], and fragment-assembly-based ones such as MC-Fold [27], FARNA/FARFAR [28-30], Vfold3D [31-34], RNAComposer [35,36] and 3dRNA [37-42]. The physics-based models are generally based on various coarse-grained (CG) representations and specific force fields, and their predictions generally involve long computational time and are still only reliable for the RNAs of small size and simple topology [6,9]. The fragment-assembly-based model are generally based on the all-atom representation, and their prediction accuracy strongly relies on templates from the PDB database since the models generally prefer to use the 3D fragments with closest sequence and length to targets in addition to fragment type [43,44]. Therefore, in spite of the great progress, RNA 3D structure prediction at high resolution and at high efficiency still remains a challenging problem [7,45-50].

In this work, we proposed a new automated and efficient RNA 3D structure building model with secondary structures as input, namely FebRNA. FebRNA involves a new structure building strategy through fragment assembly based on fragment ensembles: (i) establishing a library of CG fragment ensembles from the PDB database according to fragment type and length; (ii) building 3D CG structure ensemble through fragment assembly based on fragment ensembles; (iii) identifying top-1 CG structure from 3D CG structure ensemble through a specific CG scoring function; and (iv) rebuilding the all-atom structure from top-1 CG one and performing structure refinement. The extensive examination indicates that FebRNA gives stably reliable predictions for extensive types of RNA structures, in a comparison with existing top fragment-assembly-based models.

## METERIALS AND METHODS

FebRNA for building an RNA 3D structure was briefly illustrated in Fig. 1, and is described detailly in the following.

**Figure 1.**
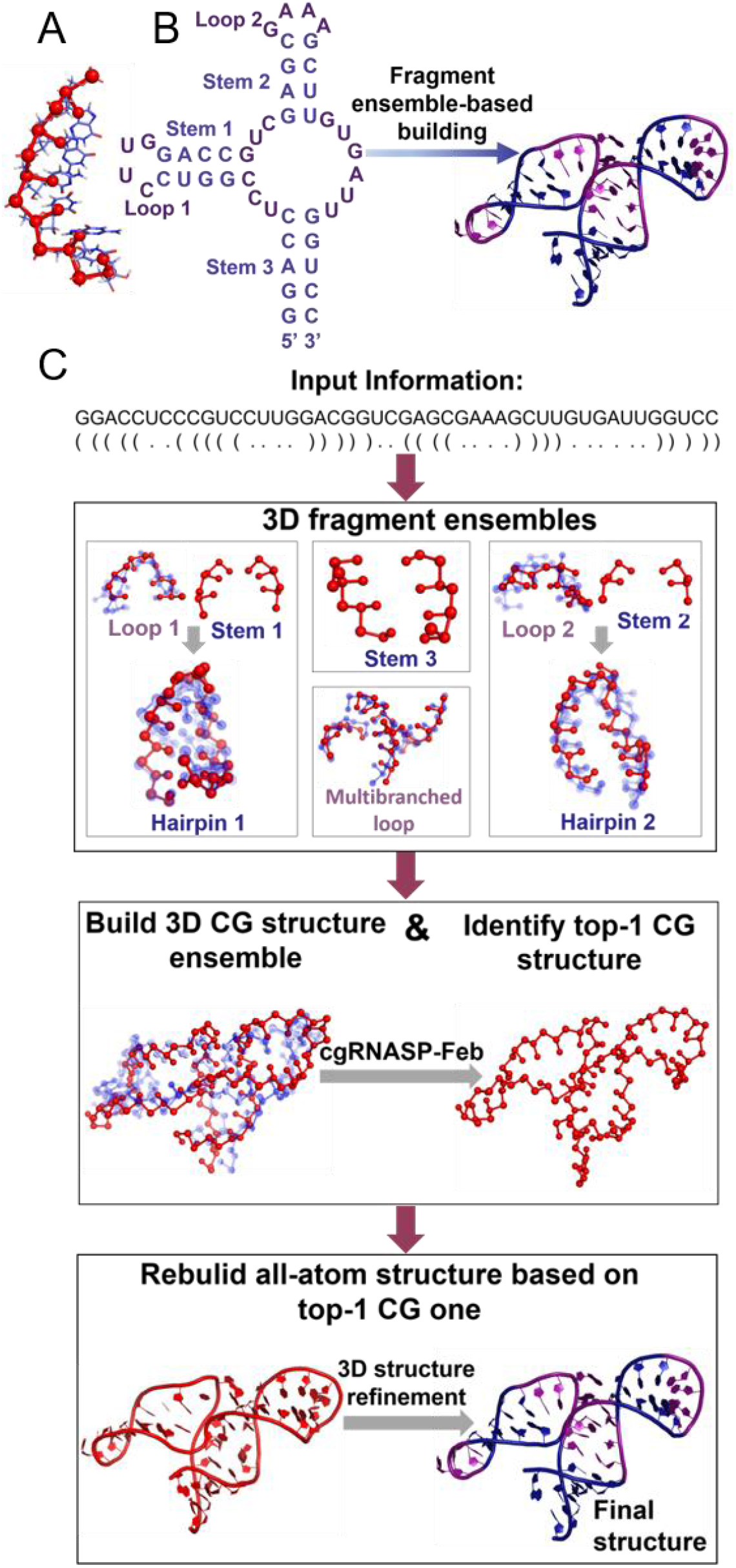
Illustration for the present model of FebRNA. (A) Coarse-grained (CG) representation of an RNA fragment in FebRNA: 3 CG beads at heavy atoms of P, C4’, and N1 for pyrimidine (or P, C4’, and N9 for purine) [15,23,31,54]; (B) From secondary structure to tertiary structure for an RNA by FebRNA. (C) Schematic of the basic steps of FebRNA. The 3D structures were displayed by Pymol [76].

### CG representation and library of 3D fragment ensembles

To simplify the library of various types of fragment structures and to improve computational efficiency in structure building and evaluation, in the library of fragment ensembles of FebRNA, each nucleotide is simplified into three CG beads at heavy atoms of P, C4’, N1 for pyrimidine (or N9 for purine) according to previous CG modeling for RNA 3D structures [15,31,51-54].

Based on the CG representation, we established the library of 3D CG fragment ensembles through the following steps. First, we downloaded the RNA 3D structures from the Protein Data Bank (PDB) (our library was updated in June 2021). Second, all 3D RNA structures were separated into basic structural fragments in the CG representation such as hairpin loop, internal loop, bulge loop, multi-branched loop, pseudoknot loop and stems, and some (1, 2 and 3) additional base pairs were reserved at each loop fragment interface which can serve as structure building entry points. Third, we classified the obtained CG fragments according to their types and lengths to build our library, and fragments of the same type and length were hierarchically clustered by a specific root-mean-square-deviation (RMSD) threshold of 1.5Å to eliminate redundancy and ensure keeping the representative fragments. If there are too many modules for a type of fragments of a length (e.g., 4-nt hairpin loop), we increased the RMSD threshold to a large value to keep the number of modules of a type and a length less than 30 for loops and less than 5 for stems, respectively. We also used Nucleic Acid Builder of AMBER [55,56] to generate canonical A-form RNA stems of different lengths to establish another library of stems for building structures of large RNAs. Finally, in the library of FebRNA, there was a non-redundant CG structure ensemble for every type of fragments of a length except for the library of canonical stems.

### Building CG 3D structure ensemble based on fragment structure ensembles

FebRNA requires the input of a sequence and its secondary structure in the dot-bracket form (for example: 5’-(((((..((((((..))))))..((((….))))……))))).-3’; PDB code, 2mtj) [57-59]. FebRNA records the stems with array subscripts from 5’ and 3’ ends in sequences and infers the types of loops and the lengths of the loops and stems. Afterwards, the dot-bracket form of a secondary structure is converted into its secondary structure tree (SST) [60] and each stem is treated as a node of the tree structure [61], which facilitates the structure building order of fragments; see Fig. S1 for the details of a typical example and Fig. S2 for building pseudoknot structures in the Supporting Information. Next, according to SST, FebRNA builds a global 3D structure ensemble based on fragment ensembles through the sequential superposition between additional (from 3 to 1 base pairs) CG atoms of a loop and CG atoms of a stem in the order of SST. Finally, FebRNA generates a CG 3D structure ensemble for an RNA.

During the building process, FebRNA automatically prefers to select the loops with more stem base pairs at their ends in order to obtain final structures at higher accuracy since more stems would generally lead to more proper orientation of loops. Furthermore, to avoid too huge numbers of final built structures for large RNAs with many interfaces, partial fragment structures in ensembles is randomly selected in building global structure for the ordinary fragments such as hairpin loops, bulge loops and internal loops except for crucially important multi-branched loops and pseudoknot loops. To obtain stable predictions, FebRNA adopts a 3-thread parallel algorithm to build a global 3D structure ensemble with up to ∼25000 CG structures for large RNAs. The present FebRNA was designed for single-stranded RNAs, and if there is ‘&’ in dot-bracket form of input secondary structures, ‘&’ was artistically treated as ‘.’ in building their 3D structures and such nucleotide of ‘.’ is removed when the structure building is done.

### Identifying top-1 CG structure through a specific scoring function

In FebRNA, to identifying top-1 CG structure from a built 3D CG structure ensemble with a huge number of candidates, we developed a specific CG scoring function of cgRNASP-Feb, according to the newly released CG statistical potential of cgRNASP [62] and our CG model for RNA folding [23-26,51,52]. cgRNASP-Feb is composed of a pure cgRNASP-like statistical potential and a bonded potential from our CG RNA folding model [23-26,51,52]. The cgRNASP-like statistical potential was derived completely in the same way as that of cgRNASP with the same distance bin width and distance cutoffs, and the bonded potential was the same as that of our CG RNA folding model [23-26,51,52]. The weight parameters in cgRNASP-Feb were optimized from a decoy training set generated from FebRNA; see the Supporting Information for the details of cgRNASP-Feb. After the structure evaluation by cgRNASP-Feb, a top-1 CG structure is identified from the built 3D CG structure ensemble with a huge number of candidates for an RNA.

### Rebuilding all-atom structure and structure refinement

In FebRNA, the top-1 CG structure by cgRNASP-Feb is converted into its all-atom one for practical use. First, we established a small library of containing 8 non-redundant all-atom structures for every type of nucleotides according to their mutual RMSDs through clustering. Second, according to the sequence of the top-1 CG structure, every all-atom nucleotide from the non-redundant ensemble is assembled to the corresponding CG one through the superposition between their corresponding (P, C4’, N9/ N1 for *i* nucleotide and P for *i*+1 nucleotide) atoms on the top-1 CG structure, and the nucleotide with the smallest RMSD on the CG atoms of the top-1 CG structure is reserved for each nucleotide. Repeat the process along the whole RNA chain, and an all-atom 3D RNA structure is built for the RNA. Finally, to remove possible steric clashes and chain breaks of the rebuilt all-atom structures, FebRNA involves a structure refinement with the method of QRNAS (https://github.com/sunandan-mukherjee/QRNAS.git) [63].

## RESULTS

To examine the performance of our FebRNA, we first built a test set consisting of randomly selected 47 RNAs including 7 simple hairpins, 7 hairpins with internal or bulge loop, 8 pseudoknots, 12 RNAs with 3-way junction, 7 RNAs with 4-way junction, and 6 RNAs with 5-way junction in the PDB database, and the RNA lengths range from 17 nt to 393 nt. This dataset with wide and well-distributed structure spectrum was called as RNA-Feb set. Furthermore, we also used single-stranded RNAs in the RNA-Puzzles dataset from CASP-like blind 3D structure prediction competition as a test set. In the following, we used the RMSD values of predicted structures from their native ones as an evaluation metrics to measure the performance of FebRNA.

The predictions from FebRNA were presented in a parallel way with those from two top existing fragment-assembly models of RNAComposer [35,36] and 3dRNA [37-42] since these two models have excellent performance in building RNA 3D structures and provide convenient and efficient webservers for users. Practically, for the two test sets, we downloaded the 3D all-atom structures from the PDB database and the 3D structures were diagnosed into their native secondary structures with the software of DSSR [57,58,64]. Afterwards, the secondary structures were input into FebRNA in the dot-bracket form as well as their sequences. Extensive examination of prediction performance was performed for FebRNA, in a comparison with RNAComposer [35,36] and 3dRNA [37-42]. Practically, for RNAComposer, we used its webserver with default setting, and for 3dRNA, we used its webserver with default setting and ‘optimize’ setting for hairpins and for RNA tertiary structures beyond hairpins respectively, since 3dRNA generally involves the structure optimization for RNA tertiary structures. Thus, we made a stricter comparison with 3dRNA beyond the assemble-only level, and for a more complete comparison, we also made additional predictions of 3dRNA with the ‘assemble’ setting for those tertiary structures.

### Performance of FebRNA for RNA-Feb set

As shown in Figs. 2A-F for the 3D structures for typical RNAs of extensive types in the RNA-Feb set, FebRNA can essentially capture the global structures for different types of RNAs, including a simple hairpin, a 2-way hairpin, a 3-way junction with 2-way branches, a 4-way junction, and a 5-way junction. The prediction performance of FebRNA for the RNA-Feb set was more quantitatively shown in Figs. 3 and 4 as well as in Tables S1-S4 in the Supporting Material.

**Figure 2.**
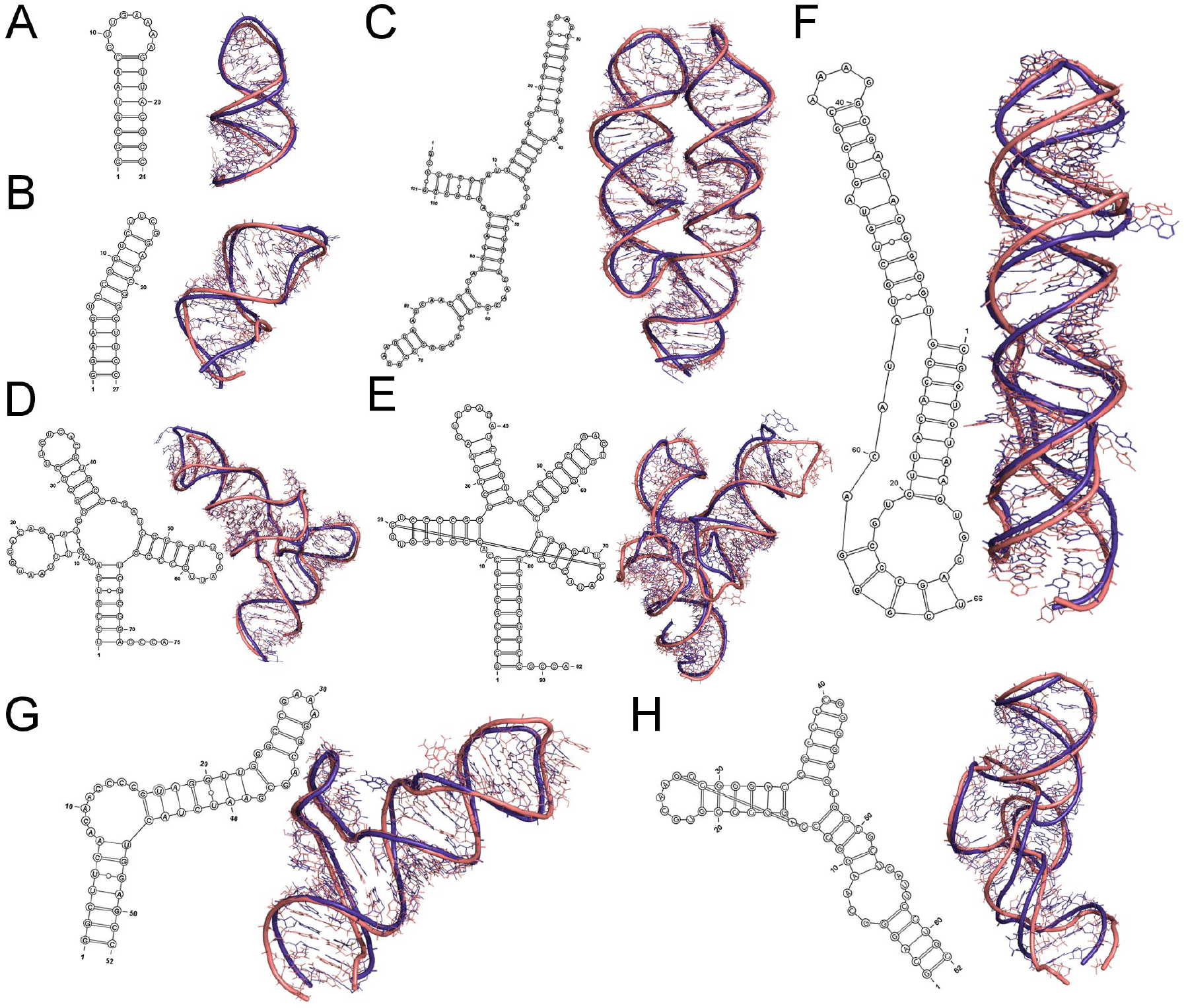
The predicted 3D structures of typical RNA structures by FebRNA: (A) a hairpin (PDB code: 1mt4); (B) A hairpin with a bulge (PDB code: 1f6x); (C) an RNA with a 3-way junction (PDB code: 1z43); (D) an RNA with a 4-way junction (PDB code: 1vtq); (E) an RNA with a 5-way junction with hairpin loop-loop contact (PDB code: 3adb); (F) an RNA pseudoknot (PDB code: 7lyj); (G) an hairpin with two 2-way junctions in the RNA-Puzzles (PDB code: 6tb7); (H) an RNA with a 3-way junction in the RNA-Puzzles (PDB code: 5t5a). The predicted top-1 structures (gold) by FebRNA are superposed on their respective native structures (blue), and the secondary structures were diagnosed from their native 3D structures with the software of DSSR [57,58,64]. The 3D structures were displayed by Pymol [76].

**Figure 3.**
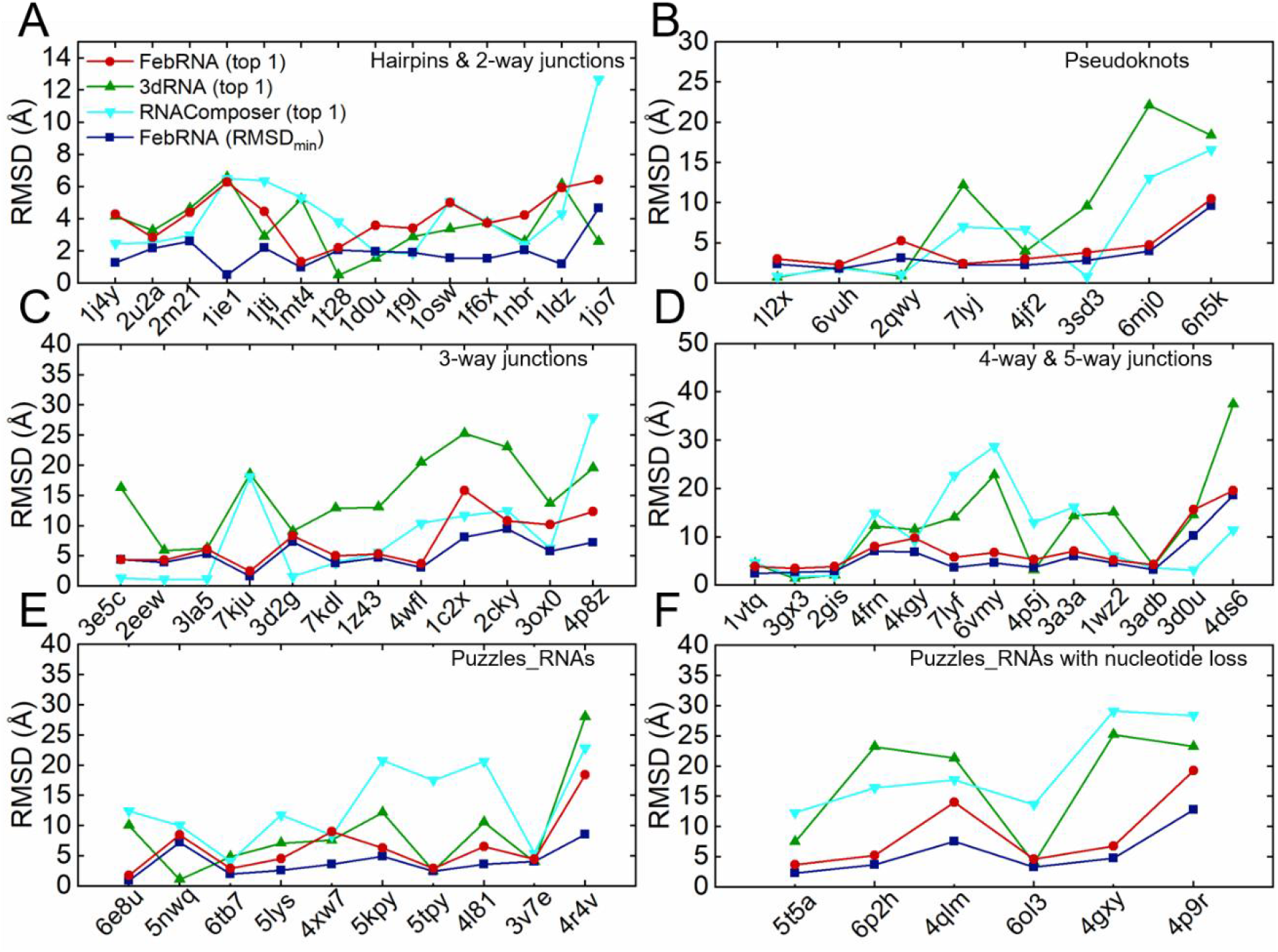
The RMSDs of the structures predicted by FebRNA for different types of RNAs: (A) hairpins without/with a 2-way junction (an internal or bulge loop); (B) pseudoknots; (C) 3-way junctions; (D) 4-way junctions and 5-way junctions; (E) single-stranded RNAs without nucleotide loss in the RNA-Puzzles; (F) single-stranded RNAs with nucleotide loss in the RNA-Puzzles. Here, RMSD_min_ denotes the RMSD of the structure closest to its native one in the structure ensemble from FebRNA, and ‘top 1’ denotes the RMSDs of top-1 structures from FebRNA, RNAComposer, and 3dRNA, respectively.

**Figure 4.**
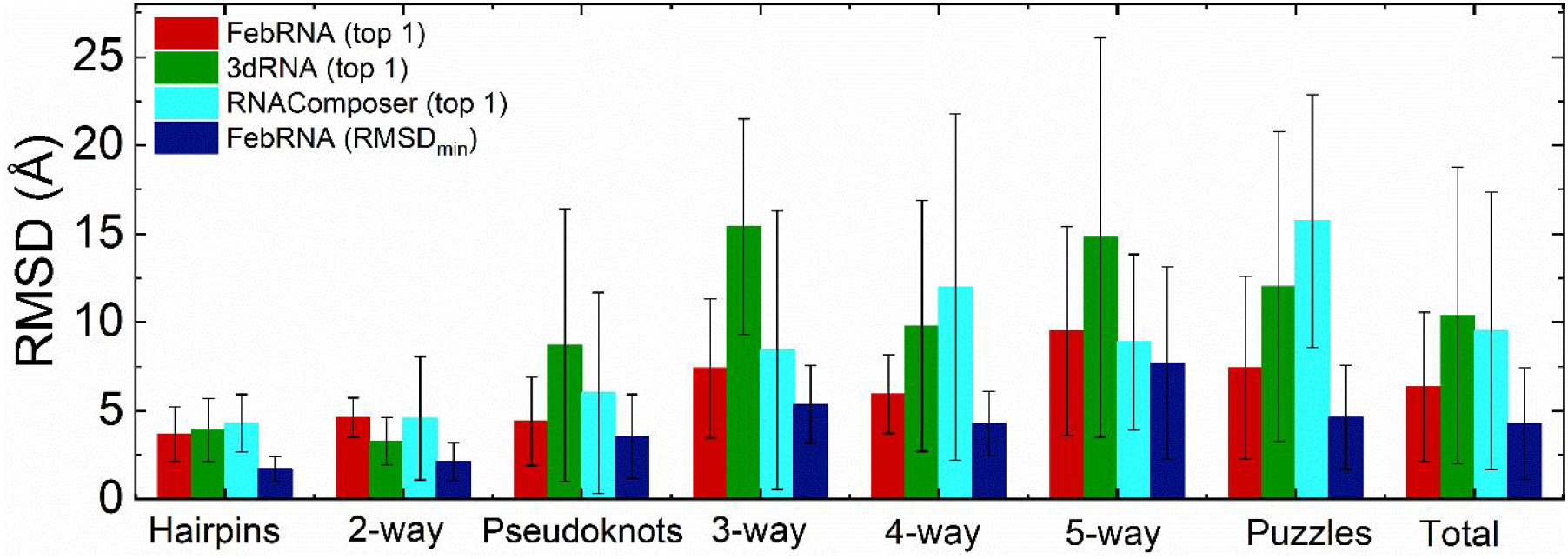
The mean RMSDs of the ‘top-1’ structures and the structures closest to native ones predicted by FebRNA for different types of RNAs, in a comparison with those from RNAComposer and 3dRNA. The error bars denote the standard deviation around the mean values.

For simple hairpins and hairpins with internal/bulge loop, the mean RMSDs of top-1 structures from FebRNA are rather small (3.7 Å and 4.6 Å), suggesting reliable predictions from FebRNA. Furthermore, such two values are overall slightly smaller than those from RNAComposer (4.3 Å and 4.6 Å), while are overall slightly larger than those from 3dRNA (3.9 Å and 3.3 Å). This indicates that FebRNA can give very slightly more accurate predictions than RNAComposer while very slightly less accurate predictions than 3dRNA for these two types of RNAs. For various simple pseudoknots with lengths between 28 nt and 127 nt, the mean RMSD of the top-1 structures from FebRNA is 4.4 Å, a visibly smaller value than those from RNAComposer (6.0 Å) and from 3dRNA (8.7 Å). Thus, FebRNA predicts 3D structures visibly closer to native ones for this kind of simple RNA pseudoknots, compared with RNAComposer and 3dRNA.

Furthermore, as shown in Figs. 3 and 4 and Table S3-S4 in the Supporting Material, FebRNA can make reliable predictions for complicated and large multi-branched structures. Specifically, FebRNA predicts the top-1 structures with mean RMSD of 7.4 Å for the RNAs with 3-way junction, and the top-1 structures with mean RMSD of 5.9 Å for the RNAs with 4-way junction. Such two mean RMSDs are visibly smaller than those from RNAComposer (8.5 Å and 12 Å) and those from 3dRNA (15.4 Å and 9.8 Å), suggesting that FebRNA makes more reliable predictions for these two kinds of complicated and large RNAs. There are very limited 3D structures of RNAs with 5-way junction in PDB, and for 6 RNAs with 5-way junction, the mean RMSD of top-1 structures from FebRNA is 9.5 Å, and the RMSD values from RNAComposer and 3dRNA are 8.9 Å and 14.8 Å. Thus, for 5-way junctions, FebRNA makes more accurate predictions than 3dRNA while very slightly less accurate predictions than RNAComposer. Such very slightly weaker performance of FebRNA (than RNAComposer) may come from the explicit and proper involvement of kissing loop-loop fragment in RNAComposer since two large (393 nt and 161 nt) RNAs with relatively large RMSDs (PDB codes: 4ds6 and 3d0u) from FebRNA have apparently kissing loop-loop interactions.

Overall, our FebRNA makes the predictions with mean RMSD of 6.0 Å for the RNA-Feb set, which is smaller than those from RNAComposer (7.4 Å) and 3dRNA (9.8 Å). This shows the overall good performance of FebRNA for the extensive RNAs with wide and well-distributed structure spectrum. Here, it is noted that the above shown predictions from 3dRNA were from the predictions of the webserver with ‘optimize’ setting [41], and the predictions from 3dRNA with ‘assemble’ setting were also shown in Tables S2-4 in the Supporting Information. It can be seen that such optimization can generally improve the predictions of 3dRNA on RNA tertiary structures especially for pseudoknots and 5-way junctions, in spite of longer computation-time cost [39], and thus the present FebRNA would become relatively more reliable at the assembly-only level. Additionally, the mean minimum RMSD of the predictions from FebRNA for the RNA-Feb dataset is rather small (∼4.3 Å) for the RNA-Feb set, which suggest that a more reliable scoring function would definitely improve the prediction accuracy of FebRNA.

### Performance of FebRNA for RNA-Puzzles set

Since the present version of FebRNA was developed for building 3D structures of single-stranded RNAs, in the following, FebRNA was examined against the single-stranded RNAs in the RNA-Puzzles dataset [65-68]. These single-stranded RNAs were divided into two categories: those without nucleotide loss and those with single nucleotide loss, i.e., there is no ‘&’ for the former and there is ‘&’ for the latter in the dot-bracket form of their secondary structures. The nucleotide loss in an RNA structure may come from the unidentifiability for the nucleotide by the experimental measurements, and the diagnosis on the 3D structure would produce a ‘&’ at the position of nucleotide loss. As shown in Figs. 2GH, FebRNA can well capture the global structures for typical RNAs in the RNA-Puzzles set including a hairpin with two 2-way junctions and an RNA with 3-way junction.

As shown Figs. 3E and Table S5 in the Supporting Information, for 10 RNAs without nucleotide loss, the mean RMSD of the top-1 structures from FebRNA is ∼ 6.5 Å, indicating the reliable predictions of FebRNA. Moreover, such value is smaller than those from RNAcomposer (∼13.4 Å) and 3dRNA (∼8.8 Å). Thus, for these RNAs in the Puzzles dataset, FebRNA predicts the structures closer to their native ones, in a comparison with RNAComposer and 3dRNA. As shown in Fig. 3F and Table S5 in the Supporting Information, for the RNAs with nucleotide loss, the mean RMSD of the top-1 from FebRNA is ∼8.9 Å, indicating acceptable predictions of FebRNA for these RNAs in the RNA-Puzzles set. Since the webservers of RNAComposer and 3dRNA cannot identify ‘&’ in the dot-bracket form of secondary structures, we performed the similar treatment to FebRNA for RNAComposer and 3dRNA, i.e., ‘&’ was replaced ‘.’ in structure building and the corresponding nucleotide was removed after the structure building process was done. With such treatment, the mean RMSDs of the top-1 structures from RNAComposer and 3dRNA are ∼19.6 Å and 17.4 Å, visibly larger values than that from FebRNA, possibly suggesting more applicable predictions by FebRNA for this kind of RNAs in the Puzzles set. Additionally, as shown in Table S5 in the Supporting Information, the structure optimization visibly improves the prediction accuracy of 3dRNA for the RNA-Puzzles set, and thus our FebRNA would appear relatively more reliable than 3dRNA and RNAComposer at the assemble-only level.

Overall, for all single-stranded RNAs in the RNA-Puzzles set, the top-1 structures predicted from FebRNA are with the mean RMSD of 7.4 Å; see also Fig. 4. Therefore, FebRNA makes consistently reliable predictions for these complex RNA structures over the wide length range from 37 nt to 189 nt. Moreover, the mean minimum RMSD from FebRNA is ∼4.6 Å for the RNA-Puzzles set, and thus the involvement of a scoring function with higher performance would further improve the predictions of FebRNA.

### Prediction efficiency and stability of FebRNA

As shown above, the present FebRNA can give consistently reliable predictions for extensive RNAs with wide and well-distributed structure spectrum and length range. It is also important to examine the prediction efficiency of FebRNA for RNAs with different lengths. As shown in Fig. 5A, FebRNA appears rather efficient in building RNA 3D structure over the wide range of RNA length. For example, for most of the RNAs, the computation time of FebRNA on a PC machine of Intel(R) Core(TM) i7-6700K CPU @ 4.00GHz CPU is less than 5 minutes, and even for the RNA with 393 nt (PDB code: 4ds6), it costs only ∼6 minutes for FebRNA to predict its structure with RMSD of 19.6 Å. Very importantly, the computation time of FebRNA appears approximately linear to RNA length, thus FebRNA has strong potential to be extended for building 3D structures of larger RNAs. The high efficiency of FebRNA may be attributed to the CG representation of RNA structures involved in FebRNA in spite of building RNA structure ensemble with a huge number of candidates. First, the CG representation would lead to more efficient superposition between two fragments during building them together; Second, the CG representation and the related removal of redundant fragments in the library would greatly reduce the searching time for an appropriate fragment; Third, the specific CG scoring function of cgRNASP-Feb can be roughly ∼100 more efficient than an all-atom statistical potential/scoring function [69-73], which leads to a very quick structure evaluation even for a huge number of candidates.

**Figure 5.**
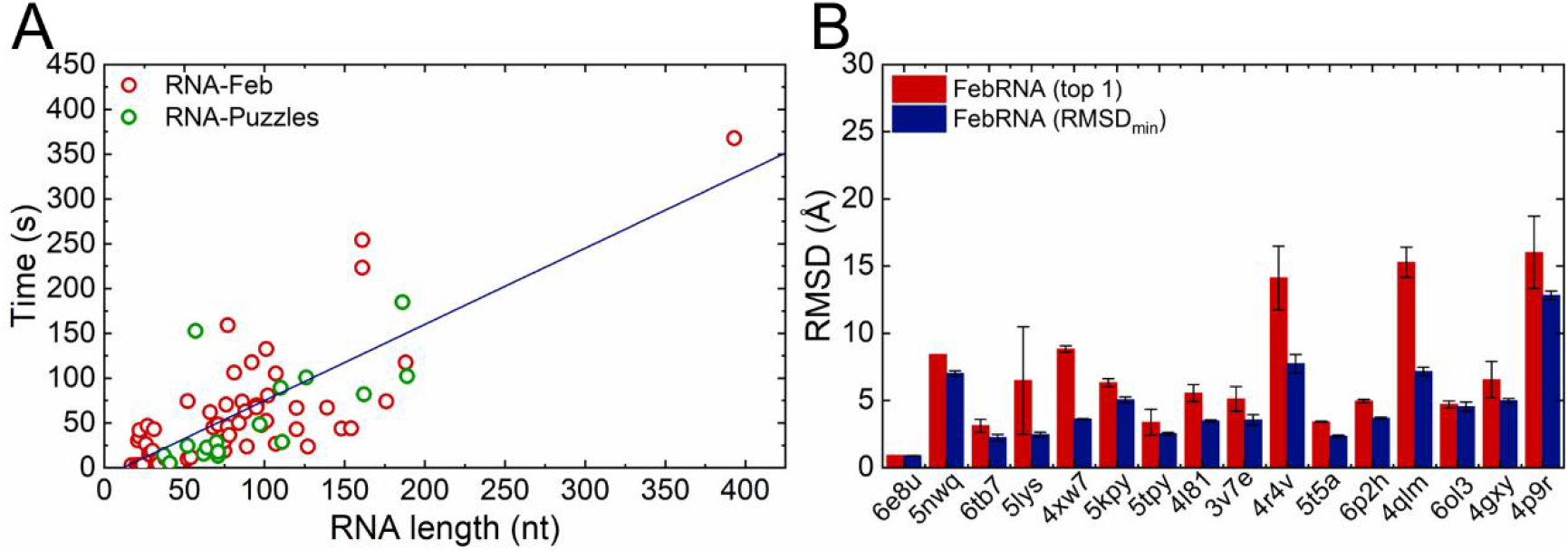
(A) The computation time (in second) of FebRNA for the RNAs in the RNA-Feb and RNA-Puzzles on a PC machine with Intel(R) Core(TM) i7-6700K CPU @ 4.00GHz CPU. (B) The mean RMSDs of the ‘top-1’ structures and the structures closest to native ones from 15 times predictions of FebRNA for the RNAs in the RNA-Puzzles, and the error bars denote the standard deviation around the mean values.

Furthermore, we examined the prediction stability of FebRNA, since FebRNA may randomly involve partial structures for ordinary fragments during building large RNA structures with many interfaces (except for multi-branched/pseudoknot loops); see Materials and Methods. We repeated the predictions for the RNAs in the Puzzles datasets 15 times, and calculated the mean RMSDs of top-1 structures and the standard deviations around the mean RMSDs. As shown in Fig. 5B, the predictions of FebRNA are rather stable for the RNAs in the RNA-Puzzles except for three RNAs (PDB codes: 5lys, 4r4v, and 4p9r). Specifically, the standard deviations for most of the top-1 RNAs from FebRNA are very small (< ∼0.7 Å), while such deviations become ∼3.5 Å for 57-nt 5lys, ∼2 Å for 186-nt 4r4v and 189-nt 4p9r, respectively. Such relatively larger deviation for the three RNAs may attributed to the non-perfect performance of the scoring function of cgRNASP-Feb, since the minimum RMSDs of predicted candidate ensemble are very stable and relatively small for the three RNAs (standard deviations are less than ∼0.3 Å); see Fig. 5B. Nevertheless, the predictions from FebRNA are overall stable and reliable. The overall stable predictions of FebRNA may be attributed to the reservation of all non-redundant structures of crucially important fragments such as multi-branched/pseudoknot loops during structure building. It is also noted that more-times predictions and a scoring function with higher performance would enable more reliable and stable predictions of FebRNA.

## DISCUSSION

As shown above, FebRNA, a model of building RNA 3D structures based on fragment ensembles, can give consistently reliable predictions for extensive types of RNAs with wide length range, in a comparison with the existing top fragment-assembly models of RNAComposer and 3dRNA. An important question may arise: How can the CG-representation-based model have good performance in building RNA 3D structures?

The good performance of FebRNA may be attributed the following reasons. First, the CG representation of (P, C4’ and N1/N9) involved in FebRNA may be a most optimal CG one with two CG atoms for describing backbone and one CG atom for describing base, i.e., such CG representation does not loss much accuracy in structure and is also convenient for rebuilding all-atom structures. Second, the removal of redundant fragments in the library also does not loss much accuracy in structure since all representative fragments were still reserved according to their mutual RMSDs. Third, through building based on fragment ensembles, a global structure ensemble with a huge number of (e.g., up to ∼25000) candidates is generated and such structure ensemble would have very strong tendency of including the structures very close to native ones. Fourth, FebRNA involves a specific high-performance CG scoring function to identify the top-1 structure from the structure ensemble with a huge number of candidates, and such top-1 structure is overall close to the one with the minimum RMSD in the built CG structure ensemble. Fifth, in building global structures, FebRNA prefers to select the loops with more stem base pairs at their ends, which would be helpful for describing the orientation of loops and improving the accuracy of final structures. More specifically, the last three treatments are responsible for the good performance of FebRNA, and the first two treatments do not bring much loss in describing structure accuracy for FebRNA.

## CONCLUSION

In this work, we have developed FebRNA, a model for building RNA 3D structures based on fragment ensembles. Extensive examinations show that FebRNA is an overall reliable and efficient model for building RNA 3D structures with secondary structures as input. Different from existing fragment-assembly models, FebRNA involves the building strategy composed of the library of non-redundant fragment ensembles at a CG representation, building CG structure ensemble with a huge number of candidates, and a specific CG scoring function for identifying top-1 structures. These involvements should be beneficial to the reliability and efficiency of FebRNA.

The present model of FebRNA can be furtherly developed for more reliably building RNA 3D structures with more complex topology. First, the current version of FebRNA was developed for building 3D structures of single-stranded RNAs and FebRNA can be continuously extended for building RNA-RNA complex structures through involving more complicated SST than that of single-stranded RNAs [74]. Second, the present version of FebRNA did not explicitly involve tertiary motifs such as loop-loop kissing ones, tetra-loop receptors, and triplex fragments, and inclusion of the tertiary contacts would definitely improve the prediction accuracy of FebRNA, especially for larges RNAs with tertiary contacts. Third, the involvement of a statistical potential/scoring function with higher performance would be crucially very helpful for improving the prediction accuracy and stability from FebRNA [62]. Fourth, although FebRNA always reserves the whole structure ensemble of those crucial fragments such as multi-branched/pseudoknot loops, FebRNA still only adopts partial structures in ordinary fragments ensembles to avoiding too huge numbers of structure candidates for very large RNAs. Such inadequate treatment may be overcome through the structure screening during structure building process by a statistical potential or a scoring function. Fifth, a structure refinement for a final structure at higher resolution may be very helpful for improve the predictions from FebRNA [75]. Finally, an involvement of a reliable CG-representation-based structure optimization with CG force fields would also be very helpful for improving accuracy of FebRNA at low computation cost [39].

Nevertheless, the present FebRNA is an overall reliable and efficient model for building RNA 3D structures with secondary structures as input for extensive types of RNAs, and the model is highly extendable for larger RNAs and for higher prediction accuracy.

## DATA AVAILABILITY STATEMENT

All relevant data are within the paper and its Supporting Information files. The package of FebRNA and the related datasets are available at website https://github.com/Tan-group/FebRNA.

## SUPPORTING INFORMATION

Supplemental Material is available for this article.

## ACKNOWLEDGEMENT

We are grateful to Profs Shi-Jie Chen (University of Missouri) and Jian Zhang (Nanjing University) for valuable discussions. The numerical calculations in this work were performed on the super computing system in the Super Computing Center of Wuhan University.

## Author Contributions

ZJT, YLT, and LZ designed the research. LZ, XW, SY and YLT performed the research. ZJT, YLT, XW, and LZ analyzed the data. LZ, XW, YLT, and ZJT wrote the manuscript.

