## Supporting Information for "FebRNA: an automated fragment-ensemble-based model for building RNA 3D structures"

---

[†]These authors contributed equally to this work.

### 1. Description of the scoring function of cgRNASP-Feb

Based on the newly released coarse-grained (CG) statistical potential of cgRNASP (1) and the CG force field of our model with salt effect for modeling RNA folding (2-6), we developed a scoring function of cgRNASP-Feb to identify top-1 structures from the candidate ensembles generated from FebRNA; see the main text. cgRNASP-Feb is composed of a cgRNASP-like statistical potential and the bonded potential from our CG RNA folding model (2-6). The short- and long-ranged potentials in the statistical potential of cgRNASP-Feb are distinguished by residue separation $k$, which is defined by $k = |m - n|$, where $m$ and $n$ correspond to the observed residue sequence positions of a pair of atoms along an RNA chain (1,7). The bonded potential in cgRNASP-Feb is used to describe the connectivity for an RNA chain (2-6). Thus, the total energy for a conformation $C$ of a given sequence $S$ can be calculated by cgRNASP-Feb (1,2):

$$\Delta E(S, C) = \Delta E_{\text{short}\,(k=1)} + \alpha\Delta E_{\text{short}\,(k=2)} + \beta\Delta E_{\text{short}\,(2\leq k<5)} + \omega\Delta E_{\text{long}\,(k\geq 5)} + \gamma\Delta E_{\text{bond}}, \quad (1)$$

where $\Delta E_{\text{short}\,(k=1)}$, $\Delta E_{\text{short}\,(k=2)}$, $\Delta E_{\text{short}\,(2\leq k<5)}$ and $\Delta E_{\text{long}\,(k\geq k_0)}$ are the energies corresponding to different residue-separation ranges, respectively. $\Delta E_{\text{bond}}$ is the bonded energy. $\alpha$, $\beta$, $w$ and $\gamma$ are weight parameters to balance the contributions of short- and long-ranged interactions, and bonded energy.

In analogy to cgRNASP (1), in the average reference state was used for short-ranged energy $\Delta E_{\text{short}}$ (7), and for long-ranged energy $\Delta E_{\text{long}}$, the finite-ideal-gas reference state (8) was used since the finite-ideal-gas reference state one has relatively good performance and has been widely used in protein structure evaluation. Please see Refs (1,7) for the details of the use of the reference states for deriving the short- and long-ranged potentials, and see Refs (9,10) for the detailed description of the existing reference states. In cgRNASP-Feb, the parameters such as the distance bin width, distance cutoffs for the potentials in $k$ ranges of 1, 2, 3-4, and $k\geq 5$ were set to the same values as those of cgRNASP (1). To recognize structure candidates with possible steric conflicts generated from fragment assembly, for the situation that some atom pairs were not observed within a certain bin width, the potentials were set to the highest potential value in the whole potential range of corresponding atom pair types and when the distance between atoms pairs is less than 3 Å (mean van de Waals diameter for CG atoms), the long-ranged potentials were set to a relatively high value of 60 $k_B$T for excluding those candidates with severe steric conflicts.

Notably, to involve the chain connectivity of RNA structures generated from FebRNA, we involved the bonded energy of our CG model with salt effect for modeling RNA folding in cgRNASP-Feb (2-6), beyond a pure cgRNASP-like statistical potential. The bonded energy is composed of bond-length energy, bond-angle energy and dihedral energy, which were initially parameterized by the statistical analysis on the available 3D structures of RNAs in the PDB database (2-6). The detailed description of the bonded energy in Eq. 1 can be found in Refs (2-6). In addition, $\alpha$ , $\beta$ , $w$ and $\gamma$ are optimized from a decoy training set generated from FebRNA (https://github.com/Tan-group/FebRNA), and they were finally set to the un-special values of 1.0, 1.0, 6.0, 8.0 and 0.01, respectively. Therefore, the scoring function of cgRNASP-Feb was derived for identifying top-1 structures from structure ensembles from FebRNA.

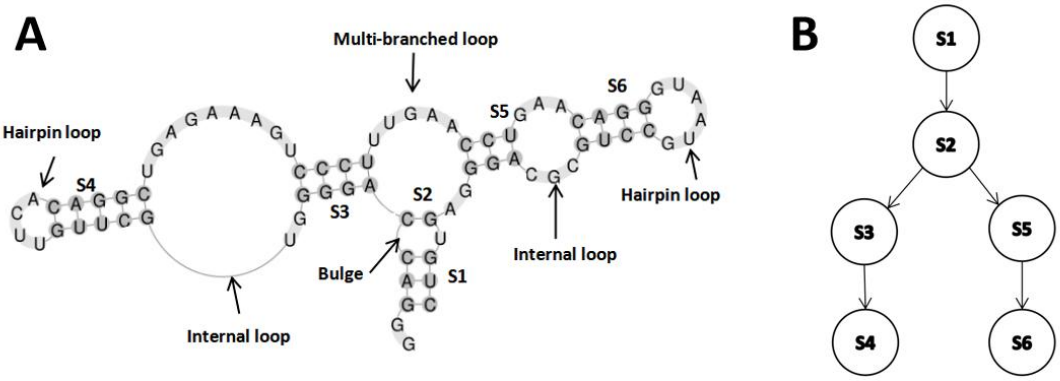


Figure S1. Illustration of the secondary structure tree (SST) for an example 75-nt RNA with three-way junction (PDB code: 3D2G). (A) The secondary structure; and (B) its secondary structure tree, where S1, S2, ..., S6 represent the nodes of the stems sequentially from 5’ and 3’ ends of the sequence.

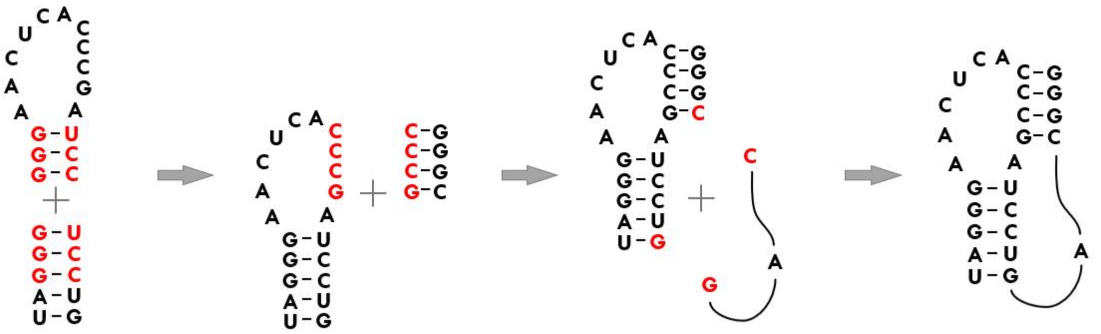

Figure S2. Schematic of building RNA pseudoknot. First, a stem and a hairpin loop are built together through the superposition between interface CG atoms (marked in red); second, another stem is built onto the hairpin loop through the superposition between side interface CG atoms (marked in red); third, another pseudoknot loop is assembled to the corresponding ends of the two stems (marked in red).

Tables S1 RMSDs of 3D structures from FebRNA for RNA hairpins without/with internal/bulge loops.

| PDB codes | Length (nt) | CG RMSD$_{min}$ (Å)[a] | CG top-1 (Å)[b] | All-atom top-1 (Å)[c] | All-atom RMSD (Å) | |
|---|---|---|---|---|---|---|
| | | | | | 3dRNA (11) | RNAComposer (12) |
| **Hairpins** | | | | | | |
| 1j4y | 17 | 1.28 | 3.77 | 4.28 | 4.17 | 2.48 |
| 2u2a | 20 | 2.17 | 2.43 | 2.85 | 3.28 | 2.51 |
| 2m21 | 21 | 2.61 | 4.08 | 4.40 | 4.66 | 3.00 |
| 1ie1 | 22 | 0.53 | 6.19 | 6.29 | 6.59 | 6.51 |
| 1jtj | 23 | 2.22 | 4.56 | 4.46 | 2.92 | 6.36 |
| 1mt4 | 24 | 1.00 | 1.00 | 1.35 | 5.22 | 5.33 |
| 1t28 | 34 | 2.07 | 2.07 | 2.20 | 0.50 | 3.82 |
| | Average | 1.70 | 3.44 | 3.69 | 3.91 | 4.29 |
| **Hairpins with bulge/internal loop** | | | | | | |
| 1d0u | 21 | 1.98 | 3.22 | 3.60 | 1.55 | 1.95 |
| 1f9l | 22 | 1.90 | 3.45 | 3.42 | 2.89 | 1.86 |
| 1osw | 22 | 1.58 | 4.42 | 5.00 | 3.37 | 5.07 |
| 1f6x | 27 | 1.53 | 3.61 | 3.74 | 3.76 | 3.80 |
| 1nbr | 29 | 2.07 | 4.07 | 4.22 | 2.62 | 2.42 |
| 1ldz | 30 | 1.19 | 6.04 | 5.93 | 6.15 | 4.31 |
| 1jo7 | 31 | 4.66 | 6.45 | 6.44 | 2.61 | 12.67 |
| | Average | 2.13 | 4.47 | 4.62 | 3.28 | 4.58 |

[a] CG RMSD$_{min}$ stands for a CG structure closest to its native one in a structure ensemble from FebRNA.

[b] CG top-1 stands for a CG structure with the lowest energy by the scoring function of cgRNASP-Feb.

[c] All-atom top-1 stands for an all-atom structure rebuilt from a CG top-1 structure.

Tables S2 RMSDs of predicted 3D structures from FebRNA for RNA pseudoknots.

| PDB codes | Length (nt) | CG $RMSD_{min}$ (Å)[a] | CG top-1 (Å)[b] | All-atom top-1 (Å)[c] | All-atom RMSD (Å) | |
|---|---|---|---|---|---|---|
| | | | | | 3dRNA (11)[d] | RNAComposer (12) |
| **Pseudoknots** | | | | | | |
| 1l2x | 28 | 2.37 | 2.68 | 3.03 | 0.76 (0.64) | 0.87 |
| 6vuh | 33 | 1.81 | 2.02 | 2.33 | 2.03 (1.83 ) | 1.88 |
| 2qwy | 52 | 3.14 | 5.14 | 5.28 | 0.90 (13.52) | 1.06 |
| 7lyj | 66 | 2.34 | 2.36 | 2.44 | 12.18 (20.84) | 7.03 |
| 4jf2 | 78 | 2.27 | 2.72 | 3.05 | 3.97 (4.53) | 6.68 |
| 3sd3 | 89 | 2.85 | 3.53 | 3.82 | 9.59 (19.96) | 0.87 |
| 6mj0 | 101 | 3.99 | 4.71 | 4.77 | 22.1 (16.61) | 13.08 |
| 6n5k | 127 | 9.62 | 10.55 | 10.48 | 18.38 (26.85) | 16.61 |
| | Average | 3.55 | 4.21 | 4.40 | 8.74 (13.10) | 6.01 |

[a] CG $RMSD_{min}$ stands for a CG structure closest to its native one in a structure ensemble from FebRNA.

[b] CG top-1 stands for a CG structure with the lowest energy by the scoring function of cgRNASP-Feb.

[c] All-atom top-1 stands for an all-atom structure rebuilt from a CG top-1 structure.

[d] The values in brackets were from the predictions of 3dRNA without optimization (only assemble).

Tables S3 RMSDs of predicted 3D structures from FebRNA for RNAs with 3-way junction.

| RNA PDB | Length (nt) | CG RMSD$_{min}$ (Å)[a] | CG top-1 (Å)[b] | All-atom top-1 (Å)[c] | All-atom RMSD (Å) | |
|---|---|---|---|---|---|---|
| | | | | | 3dRNA (11)[d] | RNAComposer (12) |
| **Three-way junctions** | | | | | | |
| 3e5c | 54 | 4.37 | 4.41 | 4.37 | 16.33 (15.10) | 1.33 |
| 2eew | 68 | 3.88 | 3.96 | 4.27 | 5.85 (5.44) | 1.09 |
| 3la5 | 71 | 5.31 | 6.15 | 6.04 | 6.25 (5.91) | 1.15 |
| 7kju | 75 | 1.61 | 2.28 | 2.47 | 18.59 (18.12) | 18.14 |
| 3d2g | 77 | 7.31 | 8.44 | 8.35 | 9.10 (20.65) | 1.59 |
| 7kd1 | 89 | 3.74 | 4.80 | 5.01 | 12.94 (23.01) | 3.82 |
| 1z43 | 101 | 4.65 | 5.19 | 5.27 | 13.08 (29.31) | 5.56 |
| 4wfl | 107 | 3.04 | 3.60 | 3.67 | 20.47 (18.23) | 10.40 |
| 1c2x | 120 | 8.10 | 15.85 | 15.80 | 25.32 (23.00) | 11.69 |
| 2cky | 154 | 9.48 | 10.70 | 10.84 | 23.04 (21.41) | 12.51 |
| 3ox0 | 176 | 5.78 | 10.18 | 10.18 | 13.71 (11.81) | 6.23 |
| 4p8z | 188 | 7.20 | 12.35 | 12.40 | 19.57 (22.24) | 27.94 |
| | Average | 5.37 | 7.32 | 7.39 | 15.35 (17.85) | 8.45 |

[a] CG RMSD$_{min}$ stands for a CG structure closest to its native one in a structure ensemble from FebRNA.

[b] CG top-1 stands for a CG structure with the lowest energy by the scoring function of cgRNASP-Feb.

[c] All-atom top-1 stands for all-atom structure rebuilt from CG top-1 structure.

[d] The values in brackets were from the predictions of 3dRNA without optimization (only assemble).

Tables S4 RMSDs of 3D structures predicted from FebRNA for RNAs with 4- or 5-way junction.

| RNA PDB | Length (nt) | CG RMSD$_{min}$ (Å)[a] | CG top-1 (Å)[b] | All-atom top-1 (Å)[c] | All-atom RMSD (Å) | |
|---|---|---|---|---|---|---|
| | | | | | 3dRNA (11)[d] | RNAComposer (12) |
| **Four-way junctions** | | | | | | |
| 1vtq | 75 | 2.34 | 3.59 | 3.84 | 4.44 (4.67) | 4.65 |
| 3gx3 | 95 | 2.71 | 3.22 | 3.49 | 1.35 (4.12) | 1.74 |
| 2gis | 95 | 2.83 | 3.43 | 3.87 | 2.06 (4.09) | 1.98 |
| 4frn | 102 | 7.05 | 7.86 | 8.02 | 12.30 (13.49) | 14.94 |
| 4kqy | 120 | 6.83 | 9.85 | 9.78 | 11.54 (12.09) | 9.10 |
| 7lyf | 139 | 3.65 | 5.70 | 5.81 | 14.01 (9.60) | 22.77 |
| 6vmy | 148 | 4.62 | 6.44 | 6.70 | 22.83 (25.68) | 28.72 |
| | Average | 4.29 | 5.73 | 5.93 | 9.79 (10.53) | 11.99 |
| **Five-way junctions** | | | | | | |
| 4p5j | 84 | 3.67 | 5.16 | 5.28 | 3.09 (3.63) | 12.96 |
| 3a3a | 86 | 5.94 | 6.76 | 7.03 | 14.37 (20.82) | 16.22 |
| 1wz2 | 88 | 4.59 | 4.83 | 5.23 | 15.09 (12.51) | 6.04 |
| 3adb | 92 | 3.16 | 3.95 | 4.23 | 4.03 (4.88) | 3.51 |
| 3d0u | 161 | 10.21 | 15.81 | 15.62 | 14.55 (25.96) | 3.08 |
| 4ds6 | 393 | 18.66 | 19.72 | 19.58 | 37.47 (44.18) | 11.47 |
| | Average | 7.71 | 9.37 | 9.50 | 14.77 (18.66) | 8.88 |

[a] CG RMSD$_{min}$ stands for a CG structure closest to its native one in a structure ensemble from FebRNA.

[b] CG top-1 stands for a CG structure with the lowest energy by the scoring function of cgRNASP-Feb.

[c] All-atom top-1 stands for all-atom structure rebuilt from CG top-1 structure.

[d] The values in brackets were from the predictions of 3dRNA without optimization (only assemble).

Tables S5 RMSDs of 3D structures from FebRNA for single-stranded RNAs without/without nucleotide loss in the Puzzles dataset.

| RNA PDB | Length (nt) | CG $RMSD_{min}$ (Å) | CG top-1 (Å) | All-atom top-1 (Å) | All-atom RMSD (Å) 3dRNA (11)[d] | All-atom RMSD (Å) RNAComposer (12) |
|---|---|---|---|---|---|---|
| **Single-stranded** | | | | | | |
| 6e8u | 37 | 0.89 | 0.93 | 1.72 | 10.02 (10.18) | 12.42 |
| 5nwq | 41 | 7.25 | 8.43 | 8.47 | 1.09 (0.26) | 10.04 |
| 6tb7 | 52 | 1.95 | 2.44 | 2.93 | 4.86 (14.03) | 4.10 |
| 5lys | 57 | 2.63 | 4.90 | 4.52 | 7.09 (11.27) | 11.76 |
| 4xw7 | 64 | 3.60 | 9.13 | 9.00 | 7.58 (10.02) | 8.37 |
| 5kpy | 71 | 4.88 | 6.17 | 6.28 | 12.17 (10.97) | 20.80 |
| 5tpy | 71 | 2.44 | 2.77 | 2.94 | 2.51 (2.46) | 17.51 |
| 4l81 | 97 | 3.60 | 6.45 | 6.57 | 10.57 (14.30) | 20.64 |
| 3v7e | 126 | 4.10 | 4.23 | 4.44 | 4.00 (4.29) | 5.43 |
| 4r4v | 186 | 8.53 | 18.46 | 18.44 | 28.00 (26.29) | 22.86 |
| | Average | 3.99 | 6.39 | 6.53 | 8.79 (10.41) | 13.39 |
| **Single-stranded with nucleotide loss** | | | | | | |
| 5t5a | 62 | 2.31 | 3.45 | 3.70 | 7.49 (15.33) | 12.31 |
| 6p2h | 70 | 3.66 | 4.93 | 5.22 | 23.23 (25.33) | 16.44 |
| 4qlm | 110 | 7.52 | 14.39 | 14.04 | 21.35 (21.82) | 17.73 |
| 6ol3 | 111 | 3.27 | 4.53 | 4.62 | 4.03 (6.66) | 13.67 |
| 4gxy | 162 | 4.80 | 6.27 | 6.75 | 25.21 (28.52) | 29.14 |
| 4p9r | 189 | 12.75 | 19.40 | 19.27 | 23.27 (33.84) | 28.39 |
| | Average | 5.72 | 8.83 | 8.93 | 17.43 (21.92) | 19.61 |

[a] CG $RMSD_{min}$ stands for a CG structure closest to its native one in a structure ensemble from FebRNA.

[b] CG top-1 stands for a CG structure with the lowest energy by the scoring function of cgRNASP-Feb.

[c] All-atom top-1 stands for an all-atom structure rebuilt from a CG top-1 structure.

[d] The values in brackets were from the predictions of 3dRNA without optimization (only assemble).
